# Transcriptome-based identification of long noncoding RNAs (lncRNAs) across the genome of *Anopheles gambiae*

**DOI:** 10.1101/2024.10.02.616138

**Authors:** Jiannong Xu, Kai Hu, Michelle M. Riehle, Vedbar S. Khadka

## Abstract

*Anopheles gambiae* is a primary malaria vector mosquito in Africa. RNA-seq based transcriptome analysis has been widely used to study gene expression underlying mosquito life traits such as development, reproduction, immunity, metabolism, and behavior. While it is well known that long non-coding RNAs (lncRNAs) are expressed ubiquitously in transcriptomes across metazoans, lncRNAs remain relatively underexplored in mosquitoes including their identity, expression profiles, and biological functions. In this study, publicly available RNA-seq datasets were leveraged to identify lncRNAs across diverse contexts, including whole mosquitoes, mosquito cells or tissues including midguts, salivary glands, and hemocytes, as well as under different physiological conditions including sugar-feeding, blood-feeding, bacterial challenges, and *Plasmodium* infections. Across this pool of transcriptomes, 2684 unique lncRNA genes, comprising 4082 transcripts, were identified. Following their identification, these lncRNA genes were integrated into the mosquito transcriptome annotation, which was then used as a reference to analyze both mRNAs and lncRNAs for transcriptional dynamics in different conditions. Like mRNAs, lncRNAs exhibited context-dependent expression patterns. Co-expression networks constructed using weighted gene co-expression network analysis (WGCNA) highlighted the interconnections among lncRNAs and mRNAs. Furthermore, we identified polysome-associated lncRNAs within polysome-captured transcripts, suggesting their involvement in translation regulation and coding capacity for micropeptides. A published ChIP-seq dataset was explored to correlate epigenetic signatures with transcriptional activities of lncRNAs. Overall, our analysis demonstrated that lncRNAs are transcribed alongside mRNAs in various biological contexts. Given their prevalence in the transcriptome, incorporating lncRNAs into transcriptome analyses will enhance our understanding of their functions, shedding light on their regulatory roles in *An. gambiae* biology.

## Introduction

Upon a biological or environmental cue, the transcriptomic response directs the production of a proteome relevant to the biological condition. Transcriptomes represent the sum of transcripts of protein-coding genes and non-coding regulatory elements controlling gene expression. To date, transcriptomic studies have been focused primarily on protein-coding transcripts. However, in recent decades, the analysis of transcriptomic data has revealed a substantial number of polyadenylated transcripts devoid of protein-coding potential across various organisms [1–3]. The evolving landscape of long non-coding RNAs (lncRNAs) represents a complex domain, only beginning to be understood in terms of their functional capacity in diverse biological processes such as transcriptional regulation, chromatin remodeling, post-transcriptional regulation, and modulation of cellular signaling pathways [4, 5]. LncRNAs have been identified in insects, including mosquitoes [6–9]. In the field of vector biology, functional genomic studies have been instrumental in elucidating the genetic basis for life traits relevant to vector competence. Recent attention has been devoted to the non-coding genome in vector mosquitoes, as highlighted in a recent review [10]. For example, genome-wide identification of lncRNAs has been conducted in both *Aedes aegypti* [11, 12] and *Aedes albopictus* [13, 14] under various biological contexts. There are 4155, 7609, and 1120 lncRNAs annotated in the reference genome of *Ae. aegypti* Liverpool, *Ae. albopictus* Foshan strain, and *Anopheles stephensi* Indian strain (UCISS2018), respectively (Vectorbase release 68). In *Anopheles gambiae*, lncRNAs have been documented in the transcriptomes from several developmental stages [15] and in the midgut following infection with *Plasmodium berghei* or *P. falciparum* [16]. In this study, we employed the transcript discovery module in the CLC Genomics Workbench to identify lncRNA transcripts across multiple transcriptomic scenarios from 12 *An. gambiae* and one *An. coluzzii* studies. These RNA-seq datasets represent diverse contexts, including distinct tissue types (e.g., whole body, midgut, salivary gland, and hemocytes), different diet types (sugar meal and blood meal), and immune challenges (bacterial and malaria infections). The identified lncRNAs were then integrated into the existing transcriptome annotation by indexing genomic coordinates for lncRNA genes. Selected transcriptomes were then used upon the updated annotation to analyze the transcriptional abundance and dynamics for both mRNA and lncRNA transcripts simultaneously.

## Material and Methods

### Datasets used in this study

We selected several RNA-seq datasets from NCBI for this study, as outlined in Table S1. The conditions of these RNA-seq studies include bacterial priming and challenge in the whole body or hemocytes; *Plasmodium berghei* and *P. falciparum* infected midgut and salivary glands, cell lines upon 20 hydroxyecdysone treatment, and sugar- or blood-fed mosquitoes as well. The RNA-seq libraries were derived from polyA-enriched RNA and sequenced with a non-stranded RNA-seq protocol.

### LncRNA identification and annotation

We developed a pipeline for the detection of lncRNA transcripts from published transcriptomes by employing the <u>module</u> Large Gap Mapping (LGM) and Transcript Discovery in CLC Genomics Workbench (v.23.0.4). The first step of the pipeline performs read mapping against the *Anopheles gambiae* PEST reference genome (v.63) with mRNA annotation. The LGM parameters allow reads to span introns, enabling the recognition of transcripts that span splice junctions. The strandedness of a transcript is determined from the splice signatures of the mapping event. The expression level of lncRNA can be low; therefore, to increase sensitivity in detecting low-abundance lncRNAs, the RNA-seq reads were pooled from a given study for the mapping step. The resulting mappings from all datasets were merged into one by the track merging tool. The merged mapping results were used as input for the Transcript Discovery module to identify transcripts. Among the identified transcripts, annotated mRNA transcripts were separated based on the annotation of PEST reference, and the remaining transcripts were examined for protein-coding potentials using the Coding Potential Calculator (https://cpc.gao-lab.org/programs/run_cpc.jsp). A transcript is classified as a lncRNA if it exceeds 200 nucleotides in length and has no coding potential. The coordinates of predicted lncRNA transcripts were annotated in the genome. The mRNA and lncRNA annotations were used as reference in RNA-seq mapping to measure the transcriptional abundance of lncRNAs and mRNAs in RNA-seq samples. The salivary gland RNA-seq dataset from Pinheiro-Sliva (BioProject PRJEB8900) was derived from *An. coluzzii. An. gambiae* and *An. coluzzii* are very closely related species [17]. The RNA-seq reads of this dataset were mapped against the *An. gambiae* PEST reference genome, so the mapped reads would be highly conserved between *An. gambiae* and *An. coluzzii*. The *An. coluzzii* specific reads would be lost during mapping as a caveat.

### Validation of lncRNA using RT-PCR

Total RNA was isolated from naïve adult female mosquitoes using Trizol following the manual. The RNA samples were treated with DNase I-XT (NEB, Catalog #M0570) to remove residual genomic DNA. To make target-specific cDNA, the reverse primer was used for priming the cDNA synthesis in each target amplicon. PCR was conducted using the SuperScript III One-Step RT-PCR system (Invitrogen, Catalog # 12574-018). The primer sequences are given in Table S2.

### Quantification of transcriptional abundance of mRNA and lncRNA

Subsequently, the updated transcriptome annotation was used to quantify the transcriptional abundance of both mRNA and lncRNA transcripts by using the RNA-seq analysis module in the CLC genomics workbench. The transcriptional abundance is represented by transcripts per million (TPM), and a false discovery rate (FDR)-adjusted p-value of < 0.05 is used to determine differentially expressed transcripts in comparison. PCA and volcano plots were used to visualize relevant expression patterns.

### LncRNA–mRNA Network Analysis using weighted gene co-expression networks analysis (WGCNA)

To gain insights into the functional attributes of lncRNAs from the co-expressed mRNA transcripts under a given experimental condition, the Weighted Gene Co-Expression Network Analysis (WGCNA) [18] method was employed to analyze the co-expression pattern of mRNA and lncRNA transcripts. The dataset from Kulkarni et al. [19] was used to demonstrate the approach. The samples in the conditions of naive, injury, *Enterobacter* challenge, and *Serratia* challenge were used. Across these samples, the transcripts with a sum TPM less than 10 were excluded from the analysis. Initially, pairwise correlations were used to identify transcripts that show similar expression patterns. Then, we determined the appropriate soft-thresholding power and calculated a signed adjacency matrix. This matrix was then transformed into a topological overlap matrix. Subsequently, we identified modules with a minimum size of 50 transcripts. The function capacity of the modules was estimated by the transcripts within the module that have gene ontology (GO) annotations.

### ChIP-seq data analysis

Gómez-Díaz et al. [20] recognized the epigenetic signatures in the midgut from the ChIP-seq data on the transcriptional active marker (H3K27ac) and the transcriptional inactive marker H3K27me3. They also identified a correlation between these epigenetic signatures and the respective transcriptional patterns of mRNAs in the midgut. We used their annotated H3K27ac and H3K27me3 signatures, and the midgut transcriptome data to examine if lncRNA expression is associated with the epigenetic signatures. The tracks of ChIP-seq H3K27ac reads, H3K27me3 reads, input control reads (without immunoprecipitation) and the midgut RNA-seq reads were aligned to visualize the transcript abundance near the H3K27ac and H3K27me3 peaks.

## Results

### I. Identification of putative lncRNAs using the Transcript Discovery approach

Similar to mRNA transcripts, lncRNA transcripts can be abundant and their expressions are modulated by experimental conditions. To identify lncRNAs expressed under various transcriptomic scenarios, we utilized RNA-seq datasets from *An. gambiae* and *An. coluzzii* available in the public domain. These datasets were derived from different mosquito body/tissue types (whole body, gut, salivary gland, and hemocytes), following different meal types (sugar-fed and blood-fed), and following immune challenges posed by bacteria and malarial parasites (*Plasmodium falciparum* and *P. berghei*). The RNA-seq datasets used in this study can be found in Table S1. To estimate the proportion of lncRNA reads in a transcriptome, we randomly selected 10 RNA-seq samples from different studies and mapped the reads against the annotated PEST genome reference. Notably, in the tested samples, 15-30% of the reads were mapped to intron and intergenic regions, indicating the presence of non-coding transcripts. Given this large fraction of mapping in non-coding regions, we employed the Transcript Discovery module in the CLC Genomics Workbench to identify lncRNA transcripts. This module facilitates the mapping of RNA-seq reads to a genomic reference with parameters allowing intron-spanning mapping. The resulting mappings were processed to infer novel transcripts. The predicted novel transcripts were examined for potential open reading frames. A transcript is defined as a lncRNA if it meets the following 3 criteria: the transcript is represented by a minimum of 10 reads across RNA-seq data sets examined, exceeds a length of 200 nucleotides, and lacks open reading frames (ORFs). Figure 1 illustrates the pipeline for lncRNA identification. The pipeline predicted 2684 unique lncRNA genes with 4082 transcripts. The predicted lncRNA transcripts are 200 – 8,672 nt in size; the number of transcripts per lncRNA gene ranges from 1-8 (97.2% of lncRNA genes have 1-3 transcripts), and the number of exons per lncRNA transcript is 1-10 (97.3% of transcripts have 1-3 exons). Overall, 59.4% of lncRNA transcripts are in the intergenic regions, 18.9% are located on the antisense strand within the gene boundaries, and 21.7% are located on the sense strands within the gene boundaries. The distribution of mRNA and lncRNA, along with their coordinates is presented in Table S3 and Figure S1. Figure 1 illustrates the lncRNA prediction pipeline, the locations of four representative lncRNAs, and seven RT-PCR validations from a cDNA sample of naive female adults at 3-5 days old. There is an EST database (expressed sequence tags, ESTdb, containing 153,332 sequences), in which the sequences represent the fragments of transcripts derived through single sequencing reactions conducted on randomly selected clones from various cDNA libraries from *An. gambiae*. Given their abundance, we predicted that lncRNA-derived fragments would be present in the ESTdb. Therefore, we did reciprocal BLASTn by searching the 4082 predicted lncRNA sequences against the EST sequences and by searching the 153,332 EST sequences against the 4082 lncRNA sequences. Using a threshold of bit score greater than 500, the BLAST revealed that 1529 EST sequences had lncRNA hits, and 611 lncRNAs had EST hits. A given EST sequence may have more than one lncRNA hit, and one lncRNA sequence may have more than one EST hit (Table S4). The presence of lncRNA hits in the ESTdb provides additional evidence for lncRNA expression.

**Figure 1.**
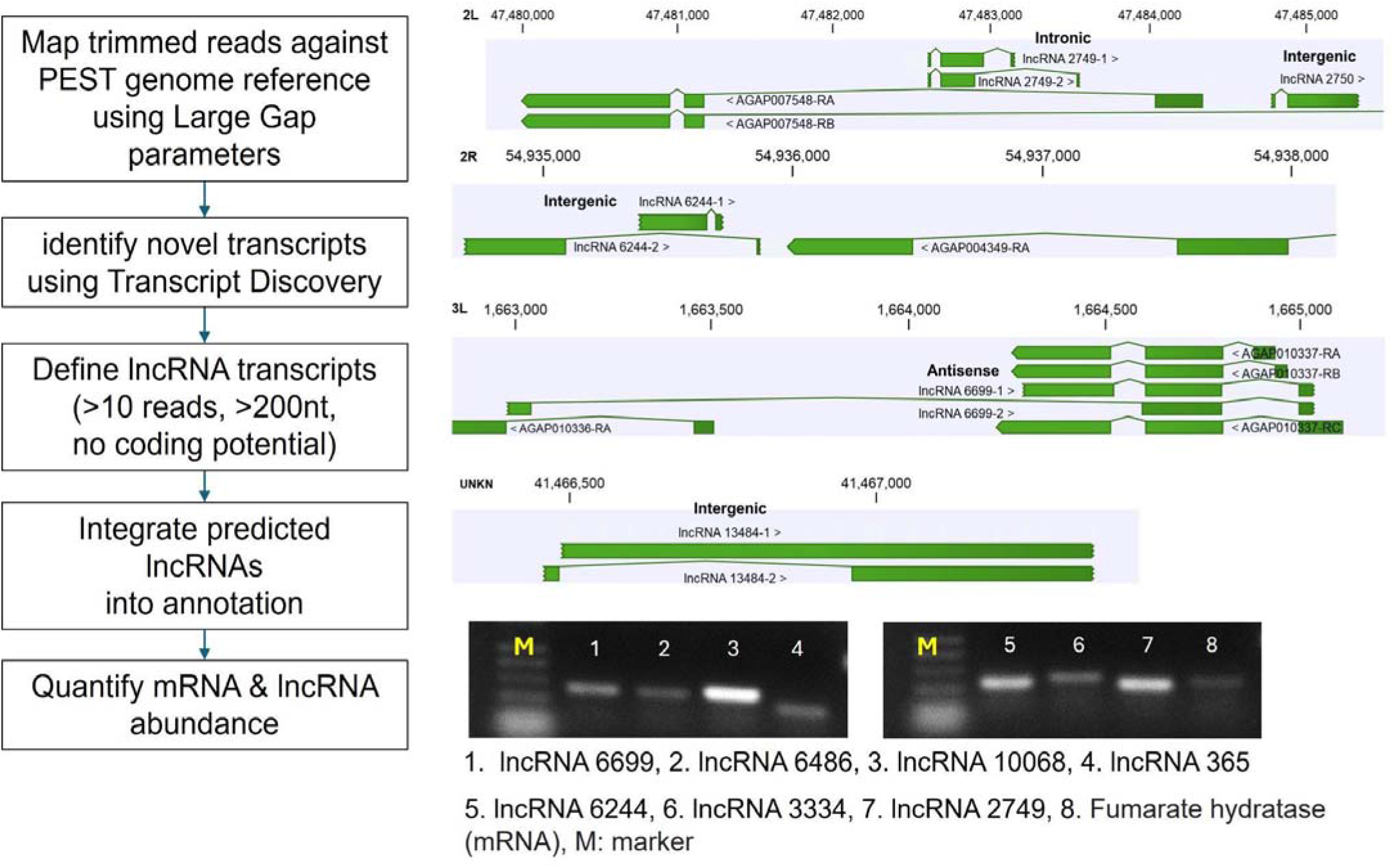
The pipeline of identifying lncRNAs from transcriptomic reads. The genomic locations of 4 exemplary lncRNAs are displayed. The gel images show the validation RT-PCR results for 7 lncRNAs and an mRNA control (AGAP001884, fumarate hydratase mRNA). M: molecular standard, 500bp, 400bp, 300bp, 200bp, 100bp.

### II. Quantification of transcriptional abundance of mRNA and lncRNA transcripts

The updated transcriptome annotation which incorporated the 4082 predicted lncRNA transcripts was used as a reference for mapping RNA-seq reads to quantify the abundance of both mRNA and lncRNA transcripts. We analyzed published datasets from two independent studies. The dataset from Kulkarni et al. (2022) has six transcriptomes representing gene expression responses to different bacterial priming regimes and bacterial challenges [19] and the dataset from Yan et al. [21] including 12 transcriptomes from circulating hemocytes (CH), heart and periosteal hemocytes (HPH) and control abdominal body (A) under conditions of naïve, injury, *E. coli*, and *S. aureus* challenges. Transcript abundance was measured in TPM.

In naïve mosquitoes with native or primed microbiota, their transcriptomes were overall similar but displayed discernable distinctions, as shown in the PCA analysis of transcriptional abundance (TPM) for both mRNAs and lncRNAs (Figure 2A and B). Upon injury or infection (*Enterobacter* or *Serratia* challenges), respective transcriptomes show distinct expression patterns, clearly separated from the naive transcriptomes with different priming regimes. In the mRNA plot (Figure 2A), the three components (PC1, PC2, and PC3) captured 59.4% of the variance, whereas in the lncRNA plot (Figure 2B), the three components explained only 32.5% of the variance, and the remaining 67.5% of variance were split into more components, with each explaining only a small portion of variance, suggesting many diverse features in the transcriptomes. The observations suggest mosquitoes differentiate between priming regimes and bacterial species for challenge, leading to distinct transcriptomic responses for both mRNAs and lncRNAs. Notably, lncRNAs exhibited greater diversity than mRNAs under the same conditions.

**Figure 2.**
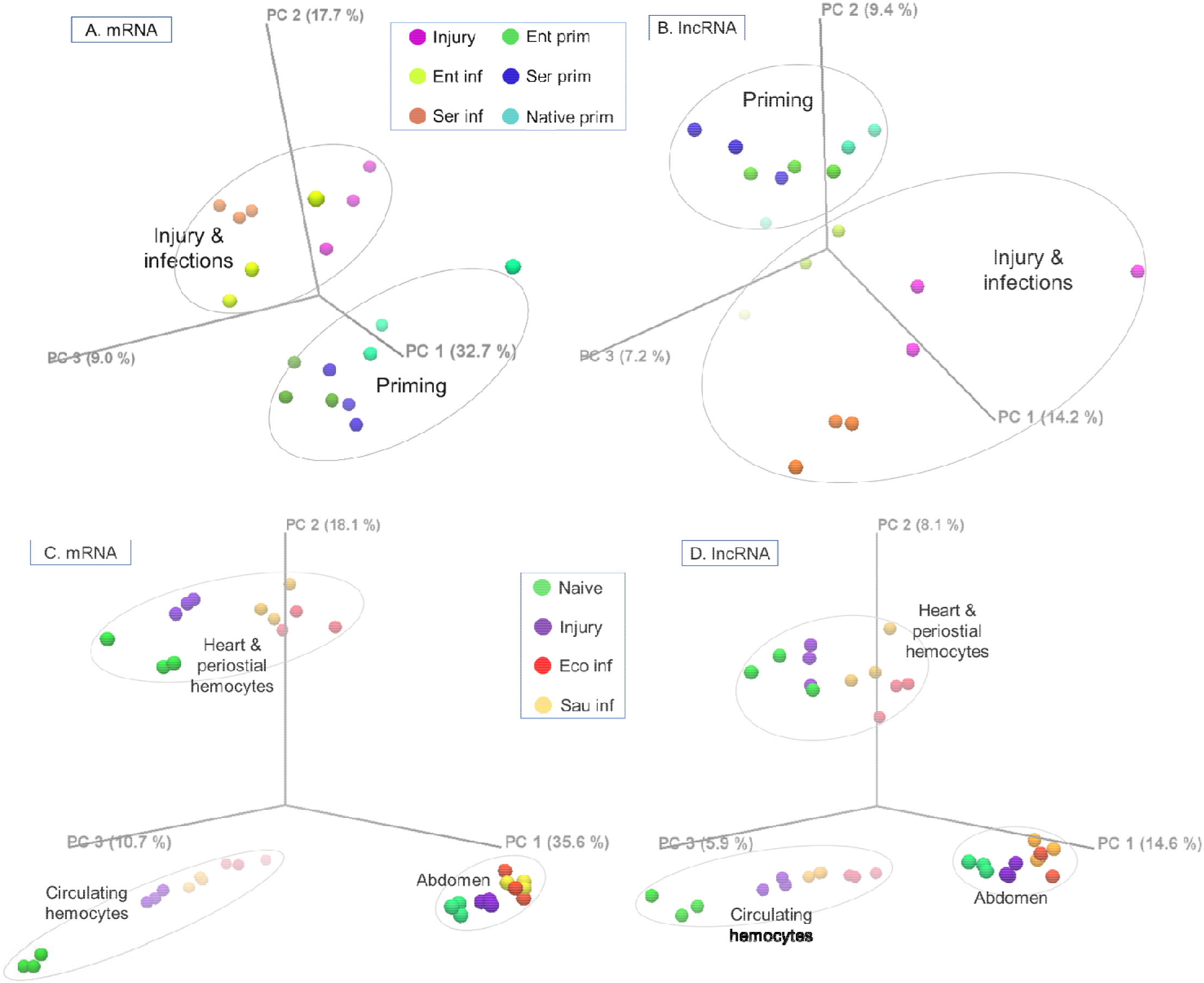
The PCA plots of TPM for mRNA and lncRNA transcripts in different conditions. The RNA-seq reads were mapped against mRNA or lncRNA annotation, and expression patterns were plotted in three principal components, PC1, PC2, and PC3. **A and B**: mRNA and lncRNA expression of whole mosquitoes upon bacterial priming and challenge. **C and D**: mRNA and lncRNA expression of circulating hemocytes, heart and periosteal hemocytes, and abdominal cells upon bacterial challenges. Ent: *Enterobacter sp*., Ser: *Serratia sp*., Eco: *E. coli*, Sau: *S. aureus*.

In Yan’s dataset [21], upon different conditions (i.e., naïve, injury, *E. coli* or *S. aureus* challenge), CH, HPH, and abdominal cells (AC) displayed distinct transcriptomic responses. Across these conditions, 8419 mRNAs and 357 lncRNAs were detected with TPM>1 in at least one condition. The difference between cell types was larger than the difference due to treatments. Both mRNAs and lncRNAs showed this pattern with a remarkable similarity (Figure 2C, 2D). In response to the treatments, the abdominal transcriptomes clustered more tightly, while CH and HPH transcriptomes exhibited greater variation, indicating that the two types of hemocytes have discrete functional assignments and play distinct roles in the responses. The first three principal components (PC1-3) of the mRNA data captured 64.4% of the total variance, whereas the PC1-3 of the lncRNAs explained only 28.6% of the variance, suggesting that lncRNAs have a more complex transcriptional diversity than mRNAs.

We conducted further analysis on the differentially expressed (DE) transcripts in different cell types, specifically AC, CH, and HPH. Figure 3 illustrates the Venn diagrams of DE transcripts (TPM>10 at least in one condition) which exhibit at least a 2-fold difference in expression and are supported by a false discovery rate (FDR) of p<0.05. Under naive conditions, pairwise comparisons revealed 2818 DE mRNAs and 78 DE lncRNAs between CH and HPH hemocytes. Additionally, we observed 3045 – 3862 DE mRNAs and 96 – 114 DE lncRNAs between abdominal cells and hemocytes (Figure 3 A, B). Upon infections, CH and HPH cells exhibited context-specific sets of DE mRNA and lncRNA transcripts. We also identified a core group of 1372 DE mRNAs and 80 DE lncRNAs that were shared across all comparisons (Figure 3 C, D). When comparing the core DE mRNAs between CH and HPH, CH showed enrichment in DE mRNA transcripts associated with energy metabolism (genes involved in glycolysis, TCA, and mitochondrial ATP production) and immune responses (Rel1, CEC2, 4 CLIPBs, 6 LRIMs, 5 PPOs). On the other hand, HPH displayed enrichment in DE mRNAs across various functional categories, including 4 transcripts of ABCC transporters, 7 transcripts in the cuticular proteins family, 16 transcripts of cytochrome P450 (CYP) genes, 12 transcripts in the GPCR, galanin/allatostatin family, and a few transcripts of immune-related genes (Rel1, lysozyme, and 2 TEP genes). Overall, the profiles of pairwise DE transcripts and the shared core DE transcripts indicate that transcriptional regulation of lncRNAs as well as mRNAs is cell-type specific and condition-dependent. These findings align with observations of lncRNAs documented in other organisms [5].

**Figure 3.**
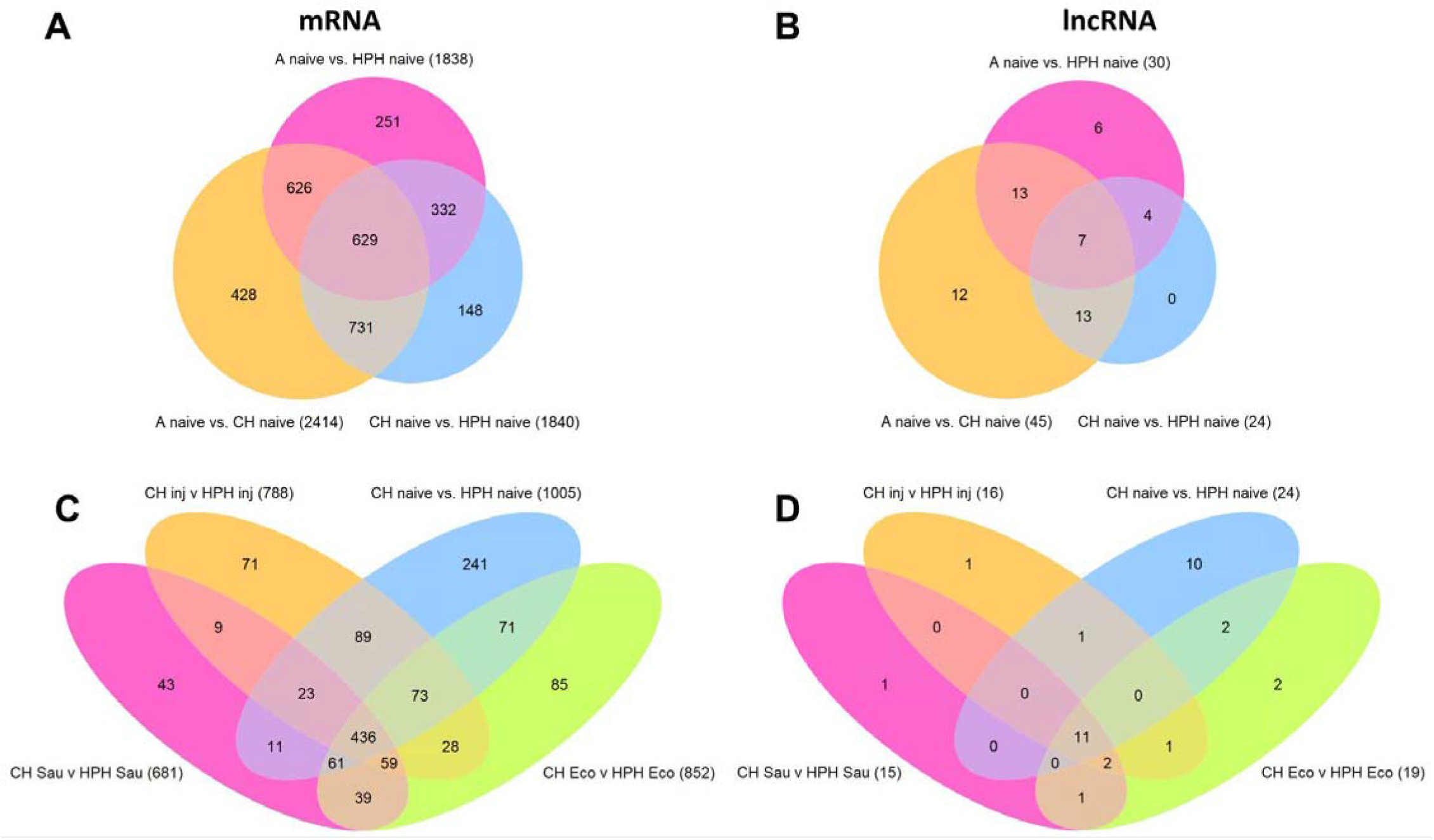
Venn diagrams of differentially expressed mRNA and lncRNA transcripts. **A and B**, DE mRNA and lncRNA transcripts in three comparisons between naive abdominal cells (A), circulating hemocytes (CH), and heart and periosteal hemocytes (HPH). **C and D**, DE mRNA and lncRNA transcripts in four comparisons between CH and HPH cells in conditions of naive, injury, *E. coli*, or *S. aureus* challenges. The comparisons include transcripts with TMP > 10 in at least one condition, a minimum absolute fold change > 2, and an FDR-adjusted p-value < 0.05.

### III. Co-expression network of lncRNAs and mRNAs

Next, we applied Weighted Correlation Network Analysis (WGCNA) to predict co-expressed networks between lncRNAs and mRNAs. To demonstrate the process, we utilized the dataset of transcriptional response to bacterial challenges from [19]. WGCNA performed transcript clustering and generated modules consisting of co-expressed transcripts. The clustering dendrogram and corresponding modules are depicted in Figure 4A. In the comparison of injury versus naïve transcriptomes, WGCNA identified 8 modules. For the *Enterobacter* challenge versus injury transcriptomes, 11 modules were identified, while the *Serratia* challenge versus injury transcriptomes yielded 18 modules (Table S5). Each module contained a mixture of mRNA and lncRNA transcripts, and the proportions of mRNAs and lncRNAs were highly correlated, as the proportion of mRNAs increased or decreased in a module, the proportion of lncRNAs tended to follow suit. The transcriptional responses to *Enterobacter* and *Serratia* infections exhibited distinct clustering/module patterns. For instance, in the transcriptome responding to *Enterobacter* infection, module turquoise was composed of 7887 transcripts (6880 mRNA and 1007 lncRNA transcripts). However, in response to *Serratia* infection, these transcripts were spread across all 18 modules. In the transcriptome upon *Serratia* infection, the dominant module turquoise contained 3412 transcripts, these transcripts were spread across 11 modules in the transcriptome responding to *Enterobacter*. This indicates that different Gram-negative bacteria induce different transcriptomic networks, which aligns well with the PCA analysis presented in Figure 2.

**Figure 4.**
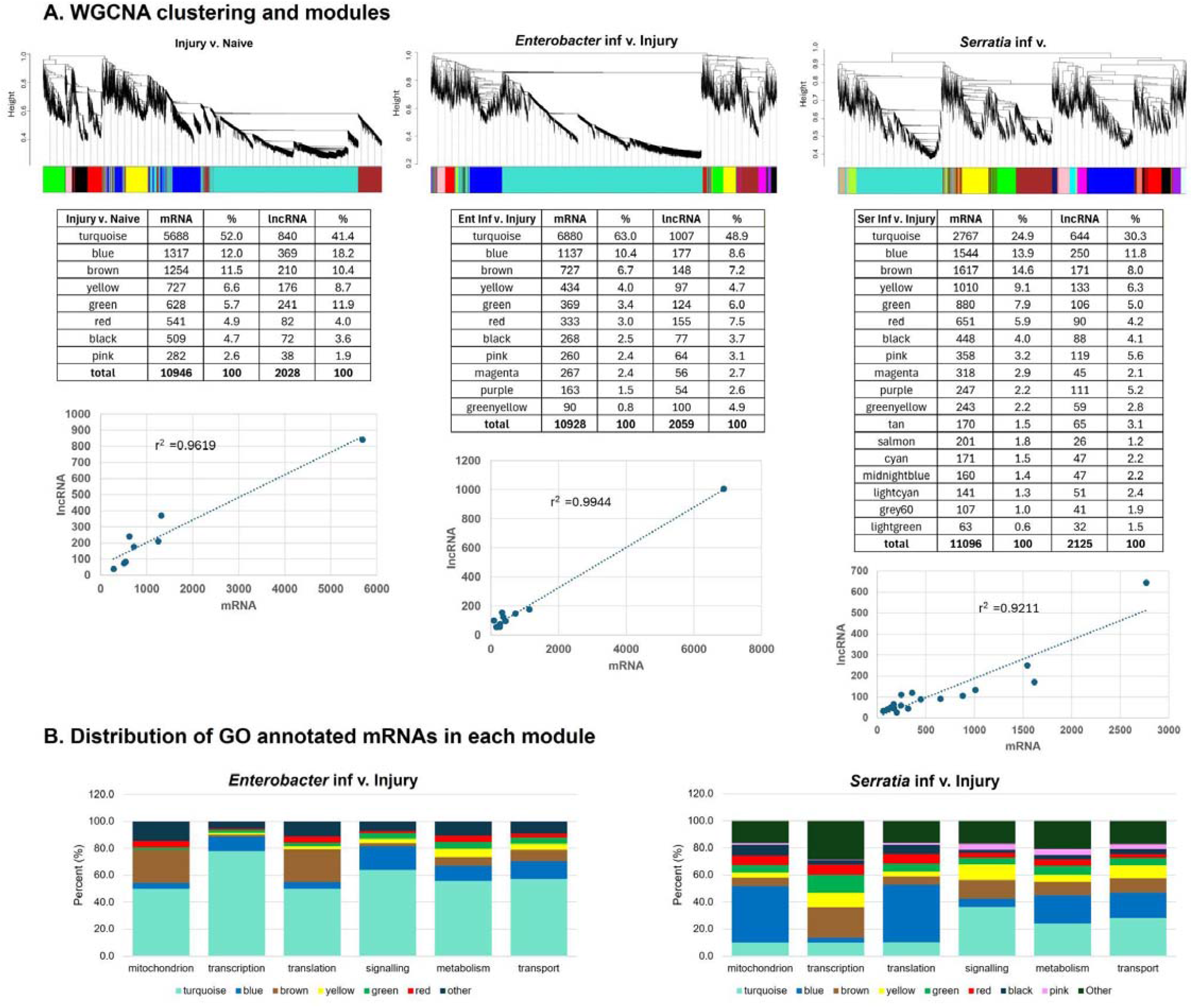
Cluster dendrogram and gene modules identified by WGCNA. **(A)** Hierarchical clustering and colored modules of co-expression of mRNAs and lncRNAs upon injury, *Enterobacter* and *Serratia* challenges. The number of mRNA and lncRNA transcripts in each module was presented in the tables under the respective cluster dendrogram. The correlation of mRNA and lncRNA counts among modules was plotted. **(B)** Distribution of modules in each of the 6 functional categories of GO annotation.

To gain insights into the potential functions of modules, we examined the gene ontology (GO) annotations for mRNAs. Approximately 70% of mRNA transcripts responsive to *Enterobacter* infection (9066 out of 12987) have GO annotations while 9464 out of 13221 (71.6%) mRNAs transcripts responded to *Serratia* infection have GO annotation. Based on their GO assignments, we categorized the mRNAs into 6 functional categories: mitochondria, transcription, translation/ribosome, signaling, transport, and metabolism. Upon the *Enterobacter* challenge, the dominant module turquoise contained mRNAs from all 6 categories (Figure 4B). For the *Serratia* infection, mRNAs from module blue were dispersed across the categories of translation, mitochondria, metabolism, and transport, while the module turquoise consisted of mRNAs from the categories of signaling, metabolism, and transport. Overall, these patterns suggest that transcriptional networks include genes from diverse functional categories. The co-expression of lncRNAs and mRNAs within these modules suggests potential functional connections including regulatory interactions between them in carrying out these functions. The correlation provides insights into the potential roles of lncRNAs based on the known functions of mRNAs within the same module.

### IV. The mRNA and lncRNA transcripts in the midgut and salivary glands post *P. falciparum* infection

To compare the expression of mRNAs and lncRNAs between the midgut and salivary glands, we utilized five datasets, which include the midgut transcriptome datasets from the naïve midguts and salivary glands of 6-to 8-day-old adult *An. gambiae* Kisumu strain [16, 20], and the *P. falciparum* Pf3D7- and Pf7G8-infected midgut 1 dpi of the *An. gambiae* L3-5 strain, the *P. berghei* infected midgut 1 dpi of the *An. gambiae* G3 strain [16], the sugar-fed naïve midgut of *An. gambiae* G3 strain [22] and the Pb-infected salivary glands 18-19 dpi of *An. coluzzii* [23]. In Ruiz et al. [24], *An. gambiae* Kisumu strain was infected with Pf3D7. The midgut transcriptome (Pf-MG) was obtained from the midgut 7 days post-infection (7 dpi) when the oocysts were encapsulated, and the salivary gland transcriptome (Pf-SG) was derived from the salivary glands 14 dpi, corresponding to the sporozoites invasion. There were three replicates for each tissue type, which enabled a statistical comparison. First, we compared the mRNAs between MG and SG and identified those that were prominently expressed in either tissue (Table S6). The Pf-MG transcriptome was enriched with mRNAs coding for protein families of aminopeptidases, carboxypeptidases, ribosomal proteins, chymotrypsins, trypsins, galectins, cytochrome P450s, aquaporins, ABC transporters, sugar transporters, and solute carrier proteins. In contrast, the Pf-SG transcriptome showed an enrichment of mRNAs encoding protein families like D7, salivary gland proteins, cytochrome P450, ABC transporters, and C-type lectins. Additionally, tissue-specific transcription factors were found in both transcriptomes. These findings were consistent with the original analyses. The TPMs from the five datasets are presented in Table S6.

It is noteworthy that immune gene sets displayed distinct patterns between the Pf-infected midgut and salivary glands. Antimicrobial peptide (AMP) genes Def1, CEC1, CEC2, CecB, and Gam1 were abundantly expressed in both tissues, but the abundance was higher in the salivary glands. Caudal, a midgut-specific transcription factor that acts as a Rel2 antagonist [25], is predominantly expressed in the midgut. Between the Pf-MG and Pf-SG transcriptomes, *Cad* exhibited 146.5-fold higher expression in Pf-MG than in Pf-SG, while *Rel2* displayed 2.3-fold higher expression in Pf-SG compared to Pf-MG (Table S6). Strikingly, families of leucine-rich repeat proteins (LRIMs), complement-like thioester-containing proteins (TEPs), prophenoloxidases (PPOs), C-type lectins (CTLs), CLIP serine proteases, and serpins (SRPNs) were expressed at higher TPM in Pf-SG than in Pf-MG, as demonstrated in Figure 5. It appears that under naive conditions, the basal expression level of these immune genes was constitutively higher in SG than in MG (Figure S2A), and Pf or Pb infection could transcriptionally affect some of these mRNAs in MG or SG (Figure S2B and S2C). The constitutive expression of these immune genes in SG was corroborated by the findings from a previous study reported by Scarpassa et al., in which the SG transcriptomes were profiled for five wild-captured Amazonian anophelines: *An. darlingi, An. braziliensis, An. marajora, An. nuneztovari*, and *An. triannulatus*. Putative proteins were predicted and annotated in the study [26]. From these annotated peptide sequences of *An. darlingi, An. braziliensis, An. marajora*, and *An. triannulatus*, the members of TEPs, LRIMs, CLIPs, and SRPNs were recognized. This transcriptional pattern of the immune genes suggests a few implications: (i) The differential expression of immune genes between MG 7 dpi and SG 14 dpi likely corresponds to the fact that the immune activities at 7 dpi have subsided in the midgut because with encapsulated oocysts that are immune-quiet, whereas the immune responses are highly active in the salivary glands at 14 dpi with ongoing sporozoite invasion; (ii) In the midgut, Caudal negatively regulates the immune gene transcription mediated by Imd pathway transcription factor *Rel2*, which may contribute to the microbial homeostasis in the midgut, as being suggested in [25]; (iii) SG is an active immune organ/tissue with a set of constitutively expressed immune genes. Intriguingly, there is an anatophysiological connection between the salivary glands and the gut. Lou et al. showed that during probing and blood feeding, the salivary gland proteins were depleted but detected by an anti-SG antibody in the midgut post a blood meal [27]. It is possible that the SG-produced immune proteins, such as TEPs, LRIMs, and CLIPs, etc., remain functional after they enter the midgut during blood feeding, which complements the Caudal controlled low production of these immune proteins in the midgut. Further investigations are needed to test this hypothesis.

**Figure 5.**
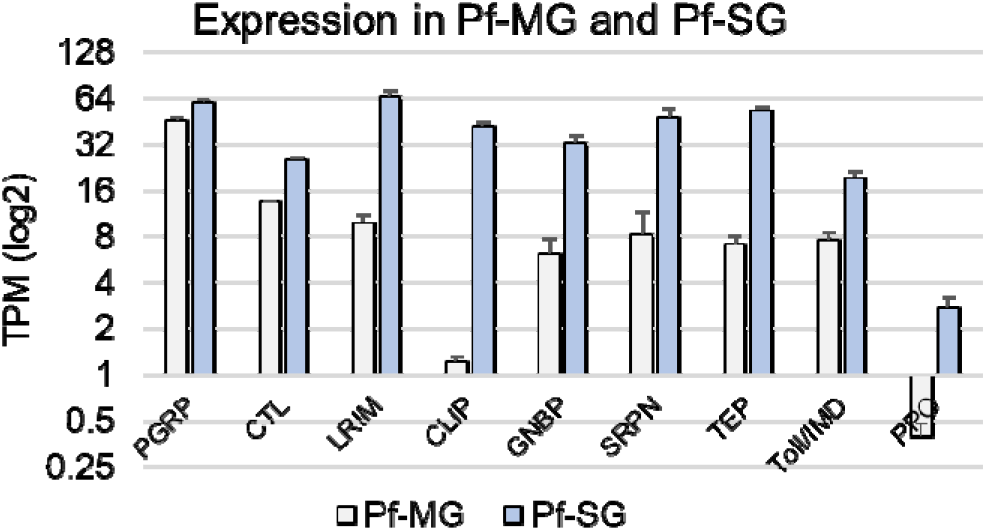
The transcriptional abundance of mRNAs in the immune categories between Pf-MG and Pf-SG. Average TPMs of the mRNAs in each immune gene category are presented. The bars represent standard error, calculated from the two replicates in the dataset.

We then analyzed differentially expressed mRNAs and lncRNAs. Using the criteria of TPM > 10 in at least one sample, FDR p-value < 0.05, and absolute fold change > 2, we identified 2596/7217 (36.0%) mRNAs and 153/501 (30.5%) lncRNAs that were differentially expressed between the Pf-MG and Pf-SG transcriptomes. The volcano plot is presented in Figure 5, and the TPM ratio of lncRNAs between the two tissues is plotted in Figure 6.

**Figure 6.**
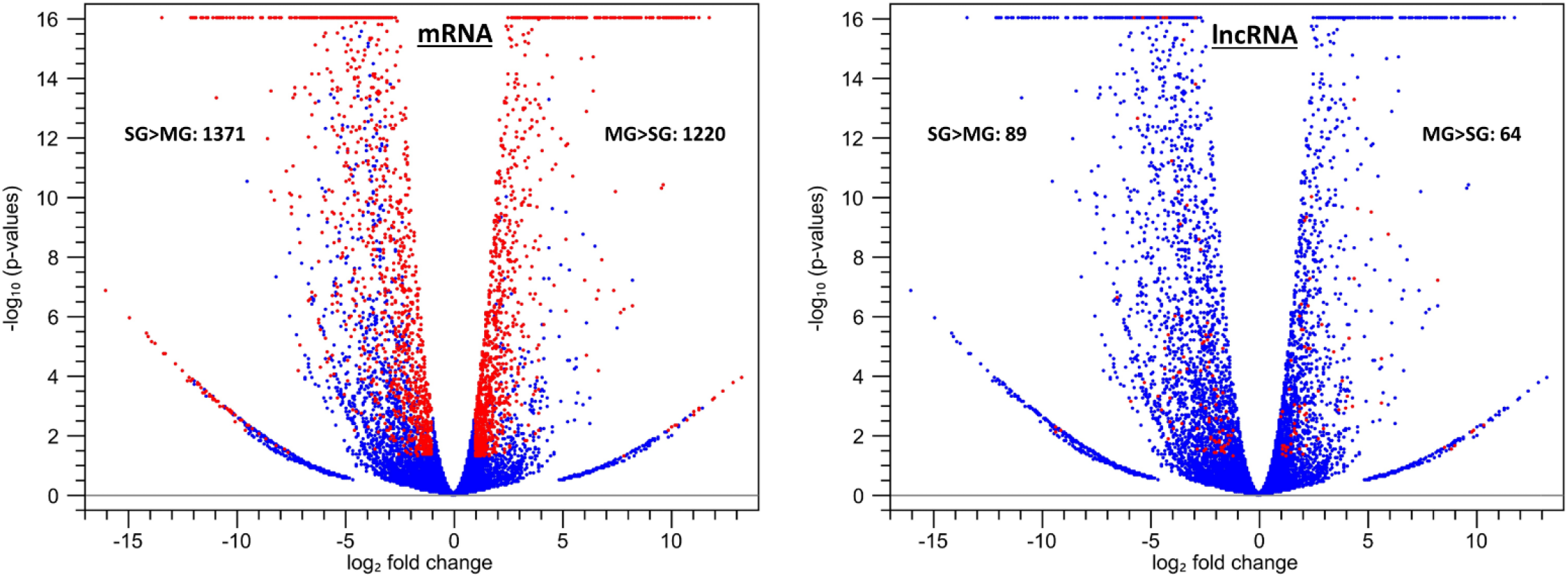
Volcano plots of differentially expressed mRNA and lncRNA transcripts between Pf-MG and Pf-SG transcriptome. The differentially expressed transcripts are highlighted in red using the filtering criteria: TPM > 10 in at least one sample, FDR p-value < 0.05, and absolute fold change > 2 between MG and SG.

### V. Polysome-associated lncRNAs

Polysome-associated long non-coding RNAs have garnered significant interest in recent years due to their pervasive presence, ability to code for noncanonical small ORFs (microproteins), and potential functional roles in various cellular processes [28–33]. In a prior investigation of mosquito transcriptomes, Mead et al. analyzed cellular and polysomal transcripts in the midgut of mosquitoes after they were fed blood meals infected with *P. falciparum* (IBM) or uninfected normal blood meal control (NBM) [34]. In this study, we used the dataset to compare the dynamics of lncRNA transcripts in the polysomal fraction (PS) and non-polysomal fraction (NP) between the IBM and NBM samples. To determine the extent of transcript association with polysomes, we calculated the polysomal portion in the total cellular transcripts as PL = PS/(PS+NP) for both the IBM and NBM samples. Like mRNAs, lncRNAs had a significant part in the polysomal portion as well. With a cutoff TPM ≥ 5, there were 314 lncRNAs in the unfractionated transcriptome, among them 251 (79.9%) were associated with polysomes. In comparison with the NBM transcriptome, IBM elevated the polysomal portion of mRNAs, which was consistent with the original analysis in Mead et al. [34]. Intriguingly, lncRNAs exhibited the same trend of polysome association as mRNAs upon the *P. falciparum* infection (Table 1). Regarding coding potential, 9 PS lncRNAs with a length range of 958-1257 nt had putative CDS for small peptides of 71-112 amino acids. These results suggest that some polysomal lncRNAs may be involved in translation regulation, and others may possess coding potential for small peptides.

**Table 1.** Polysome-associated transcripts in the IBM and NBM midgut transcriptomes*.

| PL [PS/(PS+NP)] | mRNA |  | lncRNA |  |
| --- | --- | --- | --- | --- |
|  | NBM | IBM | NBM | IBM |
|  | Count (%) | Count (%) | Count (%) | Count (%) |
| 0-30 % | 1550 (44.8) | 198 (3.5) | 27 (18.7) | 9 (3.5) |
| 31-60 % | 1481 (42.8) | 3468 (60.4) | 36 (25.0) | 79 (30.7) |
| 61-100 % | 426 (12.4) | 2075 (36.1) | 81 (56.3) | 169 (65.8) |
| Total | 3457 | 5741 | 144 | 257 |

\*The fractions of polysome-associated mRNAs and lncRNAs between NBM and IBM are significantly different, as tested by the Chi-Square test, $p < 0.0001$ .

### VI. Epigenetic signature and transcriptional abundance of lncRNAs nearby

Many lncRNAs have promoters and transcribed regions associated with active chromatin signatures, as seen in Drosophila [35], and some lncRNAs may also influence chromatin architecture [36]. Using ChIP-seq, Gómez-Díaz et al. [20] profiled the midgut (MG) epigenome for two key histone modifications: H3K27ac (associated with active promoters and enhancers) and H3K27me3 (associated with repressed regions). They also conducted RNA-seq on midguts from 6-to 8-day-old females. In this study, 6639 H3K27ac peaks and 12,939 H3K27me3 peaks were recognized in the naive midgut epigenome [20]. These two epigenetic signatures are associated with the high and low expression levels of mRNAs in the midgut, respectively [20]. We used their datasets to examine the transcriptional abundance of lncRNAs near these histone modifications. First, we mapped the midgut RNA-seq set against our mRNA-lncRNA annotation to measure the expression level (TPM) of transcripts, and then identified the histone peaks that intersect with the lncRNAs or their promoters (defined as regions located 200 bp upstream of transcription start sites (TSSs), the same criteria Gómez-Díaz et al. used in their study). According to these criteria, 307 H3K27ac peaks intersect with 243 lncRNA genes, 237 of 243 (97.5%) lncRNAs were expressed with the TPM range of 0.1-562.1 with a mean of 8.0, and 365 H3K27me3 peaks intersect with 248 lncRNA genes, and 168 of 248 (67.7%) lncRNAs were expressed with the TPM range of 0.1-14.6 with a mean of 0.9. This indicates more expressed lncRNAs were associated with the H3K27ac peaks than with the H3K27me3 peaks (Chi-Square = 75.38, p< 0.00001). This pattern suggests that lncRNA expression is also associated with both epigenetic signatures in the midgut transcriptome. Figure 7 presents four genomic regions with annotated histone modification peaks (from Gómez-Díaz et al. [20]) and nearby lncRNA and/or mRNA transcripts, along with the ChIP-seq reads mapping tracks for H3K27ac, H3K27me3, and the input background control, MG RNA-seq reads mapping tracks and respective TPM values. For instance, H3K27ac peak 1767 is associated with AGAP001548 mRNA and lncRNA 3352, both are expressed abundantly (Figure 7A), while in the region where H3K27me3 peaks 760, 315, and 202 are located, AGAP013210 and lncRNA 14014 are both expressed in a very low level with TPM less than 0.5 (Figure 7D).

**Figure 7.**
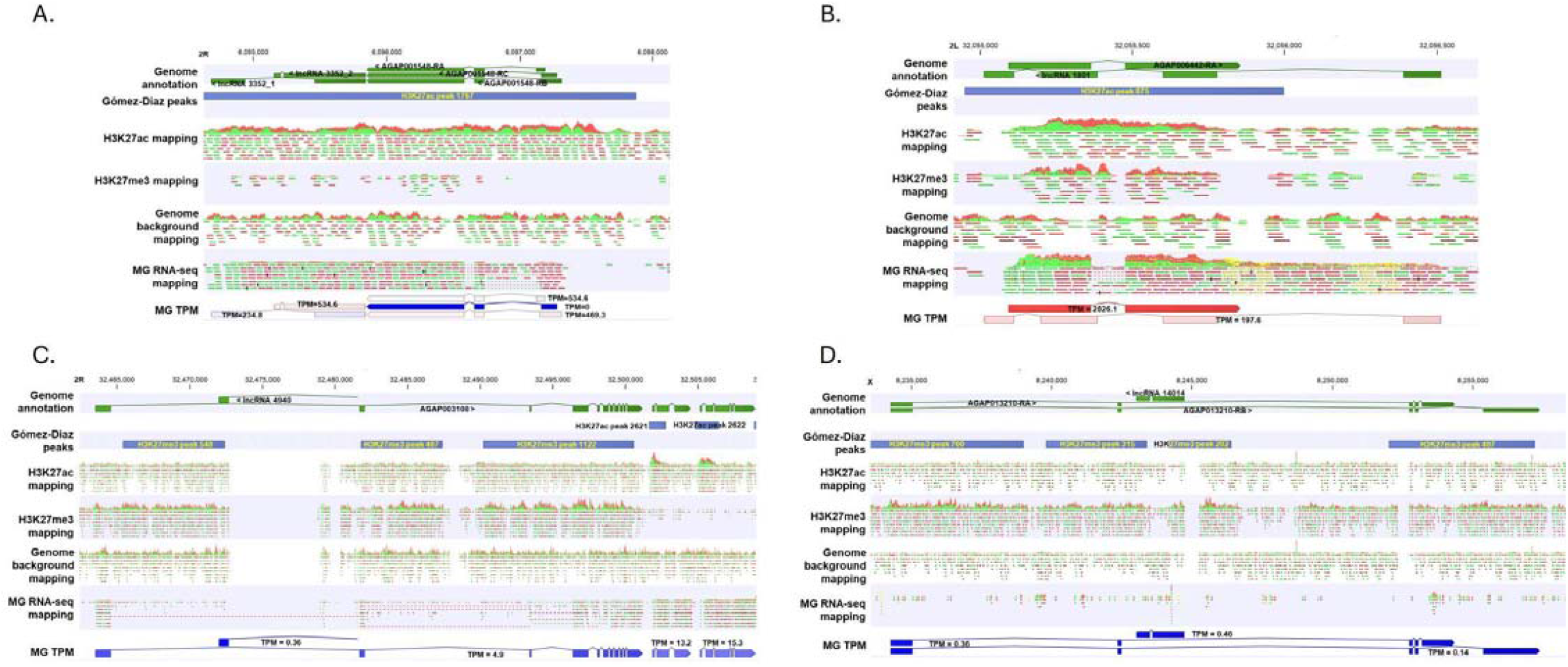
Representative genomic regions where active mark H3K27ac and repressive mark H3K27me3 and nearby genes are located. Genome annotation, annotated ChIP-seq peaks, ChIP reads mapping tracks, MG RNA-seq reads mapping and transcript TPM were aligned. Gómez-Díaz peaks were derived from the original study [20]

Enhancers are crucial DNA regulatory elements that play a significant role in initiating transcription by recruiting transcription factors [37]. Active enhancers can be transcribed into enhancer RNAs (eRNAs), which are a part of lncRNAs. Recently, Holm et al. identified 3288 active genomic enhancers from wild-caught specimens of *An. coluzzii* from Burkina Faso using Self-Transcribing Active Regulatory Region sequencing (STARR-seq) [38]. According to the enhancer coordinates, we located 278 lncRNAs on the 5’ side of enhancers and 270 lncRNAs on the 3’ side of enhancers, with a distance between 0 to 5,000 nt (Table S7). This distribution suggests possible correlations between lncRNAs and enhancers, either lncRNA expression is regulated by enhancers or certain lncRNAs represent eRNAs of co-localized enhancers. The cataloged lncRNAs described here are unlikely to include many eRNAs as eRNAs are not frequently polyadenylated.

### VII. Summary Remarks

The identification of lncRNAs from transcriptomes has been widely applied across various organisms [39–41], including mosquitoes [15, 16]. In our study, we employed a transcript discovery tool to identify lncRNAs from the selected transcriptomes derived from whole mosquitoes, midgut, salivary glands, hemocytes under different conditions (e.g., sugar-meal, blood-meal, bacterial infection, or Plasmodium infection). The predicted lncRNAs were integrated into the transcript annotation, and this updated annotation was used for transcriptome analysis encompassing both mRNAs and lncRNAs, as depicted in Figure 1. As exemplified in Figure 2, the transcription patterns of lncRNAs and mRNAs exhibited similar principal component analysis (PCA) patterns. This similarity was observed in both whole mosquito transcriptomes upon bacterial challenges and hemocyte transcriptomes upon bacterial challenges, suggesting co-expression of mRNA and lncRNA. Further validation came from the Weighted Gene Co-expression Network Analysis (WGCNA), which revealed that transcriptional networks were composed of both mRNAs and lncRNAs (Figure 4). Comparing transcriptomes from midguts and salivary glands revealed previously overlooked patterns. Notably, many immune gene families were expressed more abundantly in the salivary glands than in the midguts (Figure 5). Additionally, tissue-specific expressions of lncRNAs were identified in the comparison between the midgut and salivary glands (Figure 6). Furthermore, by mapping polysome-associated transcripts, we discovered that lncRNAs can engage with translational machinery (Table 1). Finally, we examined the genomic regions with epigenetic modifications of H3K27ac and H3K27me3, along with the expression levels of nearby lncRNAs in a midgut dataset (Figure 7). Overall, our analyses demonstrate that lncRNAs are actively expressed in all transcriptomes, and their composition and transcriptional regulation are context dependent. We recommend including lncRNAs in reference for comprehensive transcriptome analyses. The lncRNA annotation also provides a valuable resource for further investigations into the functions of lncRNAs across various life traits in mosquitoes.

## Author contributions

JX conceived and designed the study. JX, KH analyzed the data for lncRNA prediction and annotation. VSK conducted WGCNA analysis, MMR contributed to the analysis of enhancers and lncRNAs. JX wrote the original draft; all authors reviewed, edited, and approved the manuscript.

## Data accessibility

The RNA-seq datasets of the original studies are publicly available, see Table S1 for details.

## Acknowledgments

This study utilized publicly available data from the NCBI database. The authors appreciate the original research and published research articles. This work was partially supported by the Biology Department, New Mexico State University to JX. MR was supported by the National Institutes of Health, NIAID #AI145999. The authors thank Tathagata Debnath, a graduate student in the Computer Science Department of New Mexico State University, for his assistance in the data analysis in the early stage of this study.

---

**Figure S1.**
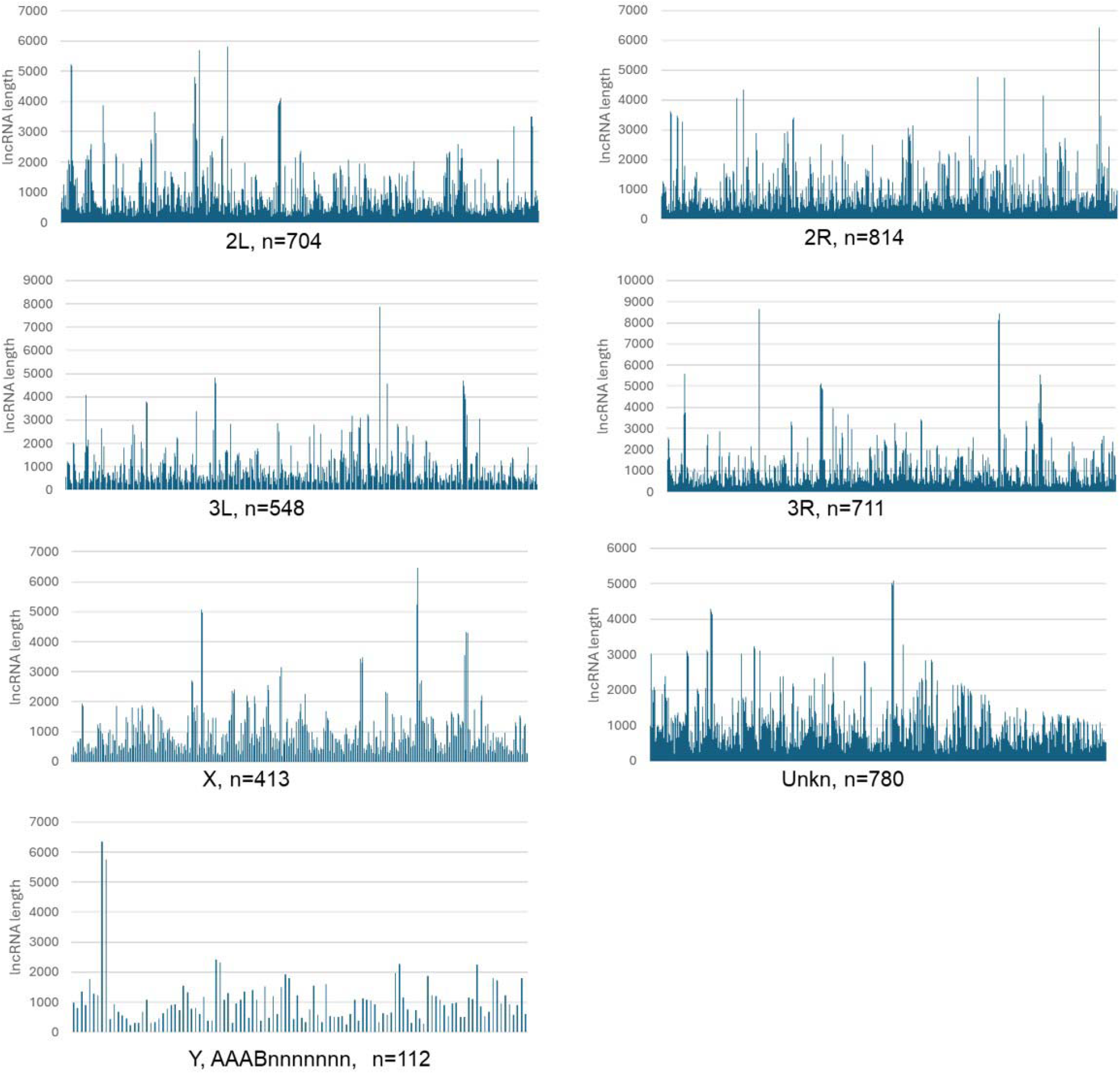
Distribution of predicted lncRNA transcripts across the chromosomes. All lncRNAs are included in the plot. The Y axis exhibits the length of lncRNA.

**Figure S2.**
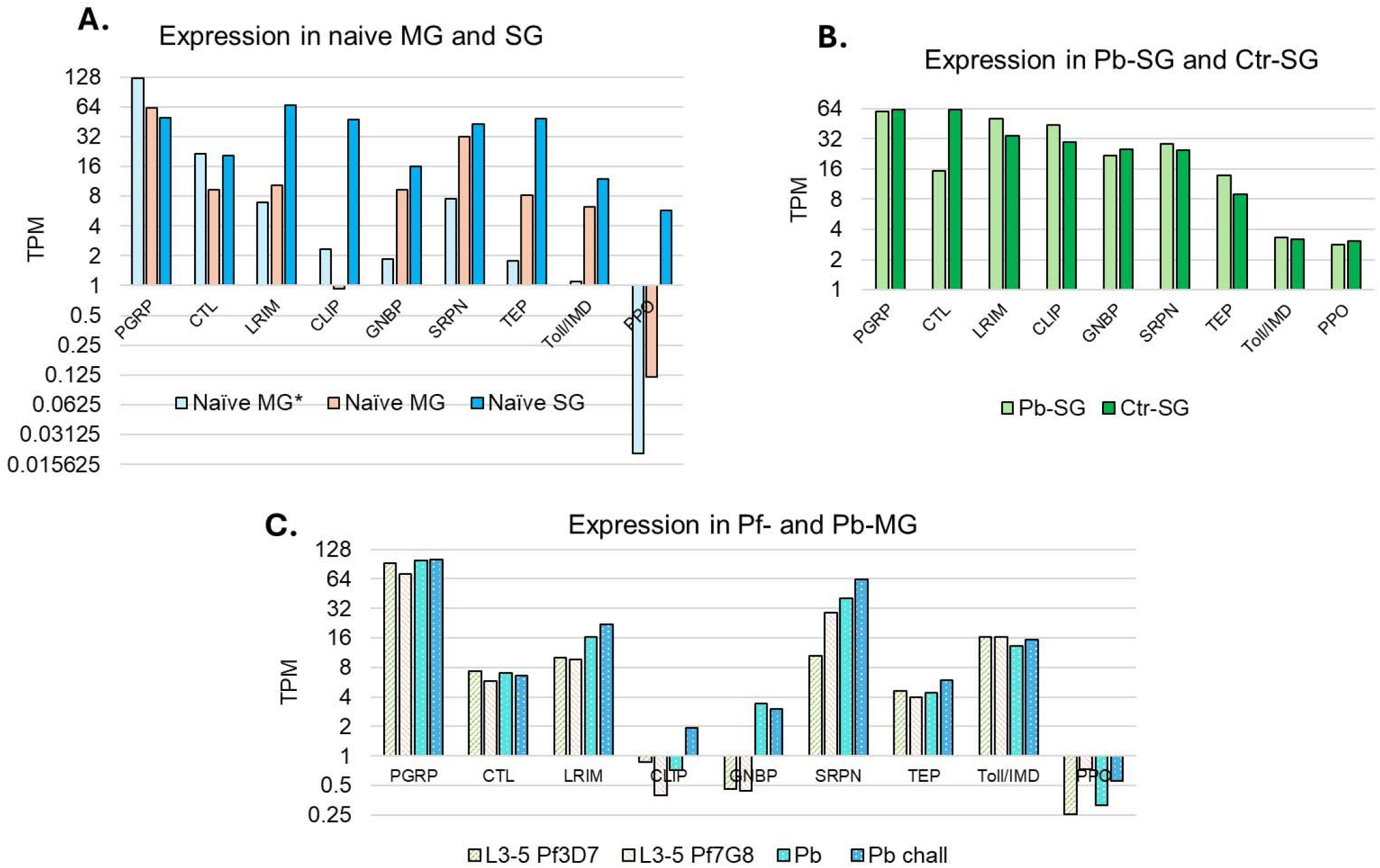
Transcriptional abundance of mRNAs in the selected immune categories. **(A)**. Expression patterns between naïve MG and naïve SG. (**B)**. Expression comparison between Pb-infected SG and control SG. (**C)**. Expression patterns in the Pf 3D7 and Pf 7G8 infected MG of Plasmodium refractory L3-5 strain and the MG with Pb infection (Pb) or Pb challenge infection (Pb chall) in G3 strain. See ref. [20] for details.

